# Polus: a Transformer-based Soft-decision Codec Enhancement Platform for DNA Storage

**DOI:** 10.64898/2025.12.16.694663

**Authors:** Lulu Ding, Kun Wang, Hongmei Zhang, Shaohui Xie, Jinlong Wang, Bo Liu, Guohua Wang, Ling Liu, Zexuan Zhu

**Affiliations:** School of Artificial Intelligence, Shenzhen University, Shenzhen, China; National Engineering Laboratory for Big Data System Computing, Shenzhen University, Shenzhen, China; College of Computer Science and Software Engineering, Shenzhen University, Shenzhen, China; Center for Bioinformatics, Faculty of Computing, Harbin Institute of Technology, Harbin, Heilongjiang 150001, China; Key Laboratory of Biological Bigdata, Ministry of Education, Harbin Institute of Technology, Harbin, Heilongjiang 150001, China; Guangzhou Institute of Technology, Xidian University, Guangzhou, China

## Abstract

DNA storage offers exceptional information density and archival longevity, but is constrained by the complex, heterogeneous errors inherent to synthesis, storage, and sequencing. Conventional error-correction schemes often rely on excessive logical redundancy to mitigate these biochemical imperfections, thereby compromising storage efficiency. Here, we introduce Polus, a deep-learning-enabled platform that bridges the gap between biochemical constraints and digital reliability through soft-decision decoding. At its core is SeqFormer, a Transformer-based channel model that synergizes sequence context with quality signals to characterize platform-specific error profiles, generating calibrated per-base confidence scores. This mechanism transforms uncertain biochemical noise into informative “soft” erasures. In *in silico* benchmarks, Polus significantly enhances mainstream codecs: it reduces the sequencing coverage required for DNA Fountain by 38.9% —increasing effective physical density by ∼80%—and eliminates persistent indel-induced errors in the Yin–Yang codec. Furthermore, it enables a targeted resequencing strategy that achieves full recovery with 99.9% less overhead than brute-force deepening. To formalize these gains and address the lack of systematic benchmarking in the field, Polus establishes a standardized nine-metric evaluation framework that rigorously quantifies the trade-offs between reliability, density, and cost. This work provides a reproducible, quantitative foundation for next-generation, context-aware DNA storage systems.

## Introduction

The exponential growth of global data—projected to reach 400 zettabytes by 2028^1^—has outstripped the capacity of conventional silicon-based storage technologies. DNA storage has emerged as a compelling alternative substrate for archival storage^2^, with unparalleled storage density (up to 215 petabytes per gram^3^), exceptional longevity^4^ (millennia under appropriate conditions), and remarkable low maintenance energy^5^. These attributes position DNA as a promising medium for next-generation archival storage, particularly for applications requiring ultra-high-density and long-term preservation, such as cultural heritage conservation and biomedical data management^6^. In a DNA storage system, digital files are encoded into oligonucleotide sequences, synthesized as DNA molecules, and later retrieved through sequencing^7^. Despite its promise, the reliability of DNA storage is constrained by biochemical imperfections across the synthesis–storage–sequencing processes^8^. During DNA synthesis (e.g., phosphoramidite chemistry), base insertions, deletions, and substitutions arise due to incomplete nucleotide coupling or depurination. Moreover, DNA strands may degrade during storage^9^ (e.g., hydrolysis-driven breaks or oxidative damage causing cytosine deamination) under suboptimal conditions. Sequencing adds further errors^10^, e.g., Illumina short-read sequencing dominates substitution error rates around 0.1–1%^11^, while Oxford Nanopore Technologies (ONT) long-read sequencing suffers high insertion/deletion errors (indels) with total rates of 1–5%^12^. These heterogeneous error sources accumulate during retrieval, necessitating robust error correction frameworks to achieve DNA’s storage potential with practical reliability.

To safeguard accurate recovery, DNA storage workflows typically employ two complementary forms of redundancy, i.e., physical redundancy and logical redundancy^13^. Physical redundancy is introduced by generating multiple molecular copies via synthesizing or PCR-amplifying many copies of the same DNA strand and sequencing high coverages. Logical redundancy encodes error resilience directly into the DNA sequences through error-correcting codes (ECCs). Early proof-of-concept studies^14,15^ established simple bit-to-base mappings, yet relying on extensive physical redundancy for recovery. Subsequent efforts introduced more sophisticated schemes by integrating ECC with biochemical constraints (e.g., balanced GC content and homopolymer-run limits). Representative examples include DNA Fountain^16^, concatenated Reed–Solomon (RS) schemes^17^, and the Yin–Yang codec (YYC)^18^. More recent designs explicitly targeted indels by coupling synchronization with ECC, e.g., HEDGES^19^, DBGPS^9^, and LDPC–watermarking approaches^20^. Nearly all these codecs correct errors with a hard-decision assumption, treating every base call as equally reliable. In reality, however, error rates vary across sequence contexts. For instance, long homopolymer runs or regions with secondary structure exhibit higher error propensities, and platform-specific error modes differ substantially between Illumina and ONT sequencing. Ignoring these variations compels designs to assume worst-case rates, thereby inflating redundancy requirements and limiting the efficiency of DNA storage.

A recent paradigm has been the advent of soft-decision decoding (SDD)^13^, which replaces the binary “correct/incorrect” treatment of each base with confidence scores, demonstrating that exploiting non-uniform base probabilities can markedly enhance decoding performance. However, the existing SDD frameworks typically assume uniform error distributions across strands and focus almost exclusively on sequencing noise, while neglecting the systematic biases from synthesis and storage. For example, in phosphoramidite synthesis the 5’-OH coupling step is markedly less efficient for guanine, leading to disproportionately high G-deletion rates^21^. Such context-dependent error patterns—arising from synthesis chemistry, storage-induced damage and platform-specific noise—remain unexploited by existing decoders.

To address these limitations, we developed Polus, an open-source modular platform with SDD for DNA storage. At its core is a Transformer^22^-based channel model, or SeqFormer for short. SeqFormer integrates multi-copy sequence information with Phred-quality scores to generate both high-accuracy consensus sequences and calibrated per-base error probabilities. Based on the probabilistic outputs of SeqFormer, Polus incorporates a codec-agnostic soft-decision decoding module to enhance error-correction capacity across representative codecs including DNA Fountain^16^, YYC^19^, and Derrick^13^. Polus also provides a standardized nine-metric evaluation framework jointly capturing density, cost, and reliability trade-offs.

Extensive benchmarking demonstrates that Polus consistently advances error identification and correction beyond existing baselines. By synergizing multiple-sequence alignment with Phred-quality signals via a Transformer architecture, SeqFormer generates calibrated per-base probabilities that enhance error enrichment by one to two orders of magnitude compared to alignment-voting methods on both Illumina and ONT platforms. When integrated with codec decoders to implement SDD, Polus significantly reduces the sequencing redundancy required for error-free recovery. Specifically, for DNA Fountain, Polus lowers the coverage threshold from 18× to 11×, thereby increasing effective physical density by ∼80% (from 1.72×10^19^ *bytes/g* to 3.09×10^19^ *bytes/g*). For YYC, Polus eliminated persistent indel-induced errors that remain uncorrectable under hard-decision decoding. Furthermore, our adaptive resequencing strategy—guided by SeqFormer’s confidence scores—targets merely ∼0.13% of low-confidence strands to achieve full recovery, reducing incremental sequencing costs by ∼99.9% compared to uniform deepening. We formalize these comparisons using a standardized nine-metric evaluation suite covering reliability, density, cost, and runtime. Collectively, these advances establish Polus as a quantitative, reproducible framework for channel modeling and context-aware decoding, providing actionable guidance for the design of high-fidelity DNA storage systems.

## Results

### Overview of Polus

Polus integrates four configurable stages namely Encoding, Simulation, Decoding, and Evaluation into an end-to-end *in-silico* DNA storage pipeline (**Fig. 1a**). Each stage is designed as an independent module with a standardized interface, allowing users to plug in different codecs or parameters and assess their impact on overall system performance. In the **Encoding** stage, digital files are firstly converted into DNA oligonucleotide libraries using a selected codec (e.g., DNA Fountain, YYC, or Derrick), producing constraint-compliant sequences with addressing and parity (**Supplementary Note 1**). Secondly, multi-source biochemical noise—synthesis errors, storage-induced damage, PCR bias and errors, and platform-specific sequencing errors—is injected during the **Simulation** stage to generate raw reads that mimic experimental outcomes for the encoded data. Thirdly, in the **Decoding** stage, reads are grouped by oligo address and processed by SeqFormer to produces each oligo’s consensus sequence along with calibrated per-base probabilities. Low-confidence positions are flagged as erasures and passed as soft inputs to the corresponding ECC layer, thereby implementing SDD. Finally, the **Evaluation** module computes a suite of nine quantitative metrics to comprehensively characterize the performance of the storage system. Unlike existing platforms that primarily aggregate codecs or simulate errors^23, 24^, Polus introduces the Decoding module which incorporates SeqFormer’s calibrated per-base probabilities to generalize SDD across codecs. This approach increases the effective error-correction capacity of existing schemes without modifying their encoders. We first validate SeqFormer’s architecture, consensus accuracy, and calibration performance, then demonstrate Polus’s consistent SDD gains with DNA Fountain, YYC, and Derrick codecs, and finally compare these codecs using the nine-metric evaluation suite.

**Fig. 1.**
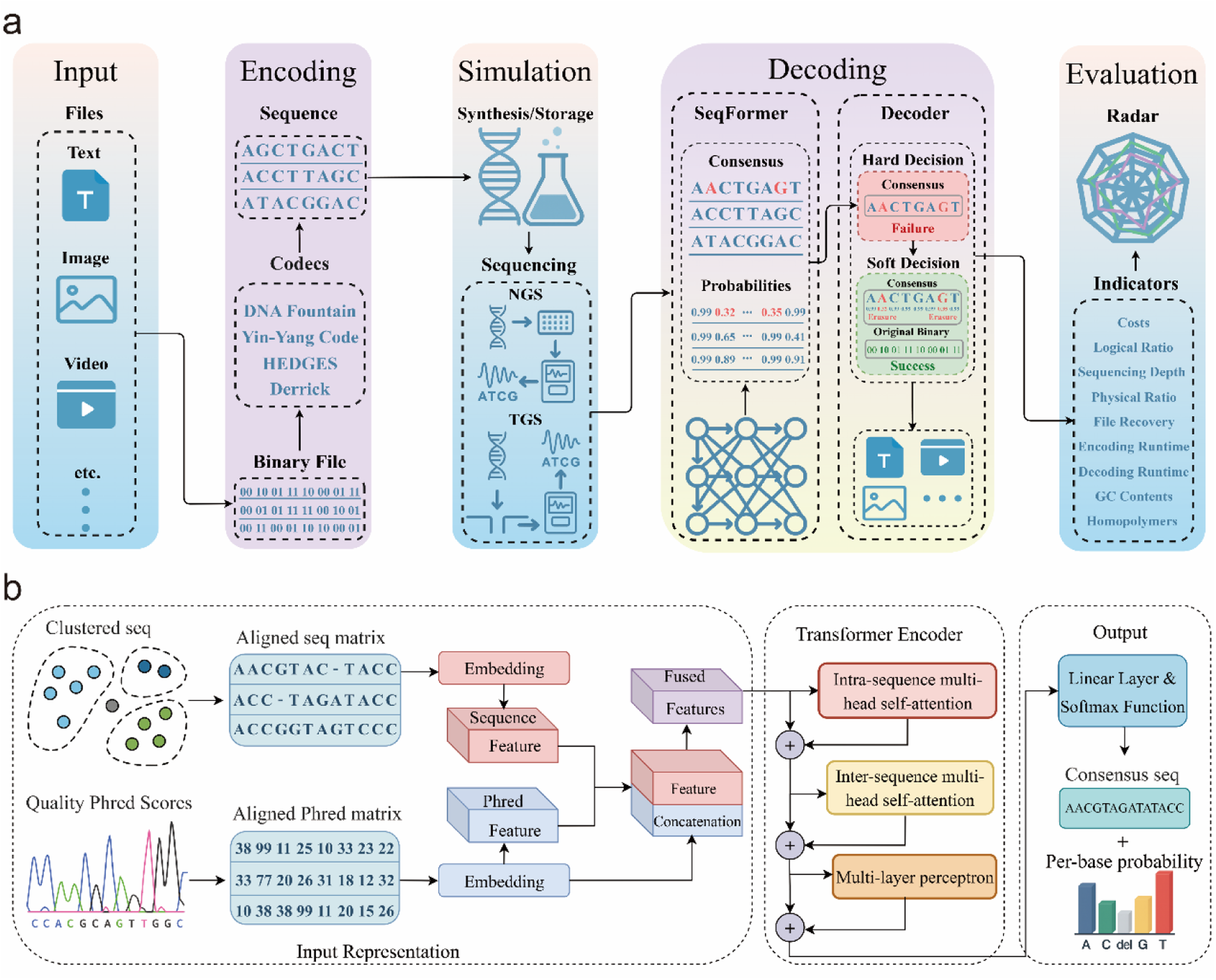
Overview of the Polus platform and architecture of the SeqFormer model. a, The end-to-end *in-silico* DNA storage workflow. The pipeline integrates four modules: Encoding (converting digital files to DNA with representative codecs), Simulation (modeling synthesis, storage, and sequencing errors), Decoding, and Evaluation. During decoding, raw sequencing reads are firstly processed by SeqFormer to generate both a consensus sequence and calibrated per-base error probabilities, which were then passed to the downstream decoder. This decoder features a dual-path mechanism: a hard-decision decoder (HDD) serves as the primary fast path, while a soft-decision decoder (SDD) is triggered upon HDD or cyclic redundancy check (CRC) failure. Low-confidence bases identified by the SeqFormer are flagged as erasures to enhance recovery, and the decoder iterates until succeeds and the reconstructed file passes the CRC. The evaluation module reports a nine-metric dashboard encompassing file recovery rate, logical and physical density, sequencing depth, runtime, cost, GC content, and homopolymer length. **b,** Schematic of the SeqFormer network. The model processes two aligned inputs: a nucleotide matrix and a corresponding Phred quality score matrix. These inputs are embedded and normalized into sequence and quality features, which are then concatenated and fused via a linear layer. The Transformer encoder comprises 8 stacked blocks containing intra-sequence multi-head self-attention, inter-sequence multi-head self-attention, and a multilayer perceptron (MLP). The model jointly outputs a consensus sequence and calibrated error probabilities over {A, C, G, T, -}, for each position, providing the soft information required for the SDD module.

### Effects of SeqFormer

#### SeqFormer architecture

SeqFormer delivers a probabilistic channel model of DNA storage at single-nucleotide resolution (**Fig. 1b**). From an alignment-synchronized pile of reads and their corresponding Phred quality scores, SeqFormer produces a per-oligo consensus sequence and a calibrated categorical distribution over {A, C, G, T, –} at each position (with “–” denoting a deletion). Concretely, multiple-sequence alignment information and the corresponding quality signals are jointly embedded and fused, per-read representations are aggregated across reads, and a Transformer encoder with dual-stream multi-head self-attention captures long-range sequence context before a lightweight prediction head yields per-base probabilities. The consensus sequence is then obtained by selecting, at each position, the nucleotide with the highest predicted probability, and the per-base error probability is defined as one minus the model’s probability for that chosen base. Architectural and training details are provided in **Methods** section.

#### Consensus accuracy

SeqFormer enables high-accuracy consensus calling with lower sequencing depth than previous methods (**Fig. 2a-b**). On the Illumina-sequenced DNA oligonucleotides from Organick *et al.*^17^ (encoding ∼200 MB data), SeqFormer reconstructed ∼95.7% of oligonucleotides error-free at only 3× coverage, slightly surpassing BSAlign^25^ and BMALA^26^ and clearly outperforming Iterative Majority, DivBMA, and Hybrid^27^ methods. By 5× coverage, all methods exceeded 98% accuracy, with SeqFormer and Iterative Majority both reaching ∼99.4%. Although Iterative Majority can slightly outperform SeqFormer at higher coverage (≥4×), its runtime grows steeply with depth, making SeqFormer far more efficient at practical coverages. Similarly, on the ONT-sequenced DNA oligonucleotides from Ding *et al.*^13^ (encoding ∼5 MB data), SeqFormer remained competitive at very low depth and became the top performer beyond ∼9× coverage, exceeding BMALA’s accuracy by 5-20% within comparable runtime (<100 s).

**Fig. 2.**
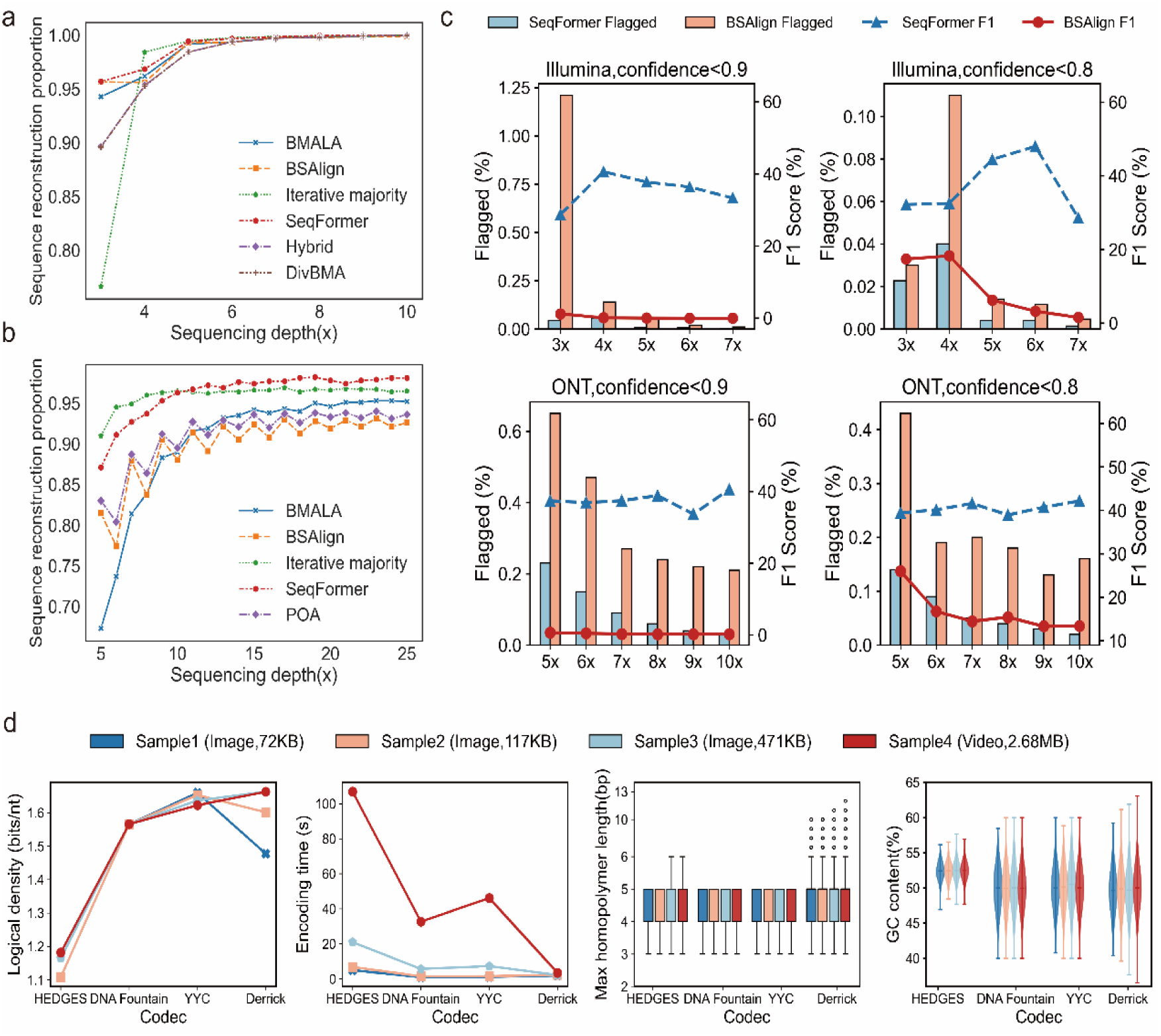
Consensus reconstruction accuracy, error enrichment validation, and codec encoding assessment. a-b, Comparison of sequence reconstruction accuracy (consensus) across varying sequencing depths for Illumina (a) and ONT (b) datasets. SeqFormer was benchmarked against established alignment methods, including Iterative majority, DivBMA (only Illumina), Hybrid (only Illumina), BSAlign, POA (only ONT), and BMALA. **c,** Performance benchmarking of error enrichment strategies on Illumina and ONT platforms under confidence thresholds of 0.9 and 0.8. The dual-axis plots contrast SeqFormer (blue) and BSAlign (red). Bars (left axis) represent the percentage of bases flagged as low-confidence (candidate erasures), while lines (right axis) indicate the F1 score (harmonic mean of precision and recall) of error detection. SeqFormer consistently achieves superior error detection, maintaining high F1 scores while flagging an orders-of-magnitude smaller fraction of bases compared to the alignment-based BSAlign. **d,** Characterization of encoding metrics for four DNA storage codecs (HEDGES, DNA Fountain, YYC, and Derrick) using four distinct file samples ranging from 72 KB to 2.68 MB. Subpanels display logical density (bits/nt), encoding time (s), and box/violin plots representing the distribution of maximum homopolymer length and GC content per encoded read.

#### Calibration of base-wise probabilities

The probabilistic outputs of SeqFormer are well-calibrated, with outputted confidence scores closely reflecting actual error probabilities (**Supplementary Figs. 1-2**). On the Illumina dataset across 3× to 7× coverage, bases in SeqFormer’s high-confidence bin (0.9–1.0) had empirical error rate dropping from 2.27×10^-4^ at 3× coverage to undetectable at 7× coverage. In contrast, the baseline BSAlign became increasingly overconfident, i.e., its high-confidence error rate increased from 1.81×10^-3^ to 4.45×10^-3^ over the same range, presumably because simple vote-based heuristics inflate confidence with read count even when systematic errors persist^28,29^. Consequently, in the high-confidence regime, SeqFormer was ≥8-fold better calibrated at 3× coverage and ≥300-fold at 6× coverage than BSAlign, with zero high-confidence errors at 7× coverage. A similar trend was observed on ONT datasets, i.e., the high-confidence error rate of SeqFormer fell by an order of magnitude as coverage increased, whereas that of BSAlign rose—implying a SeqFormer calibration advantage of ≥43-fold at 5× coverage rising to ≥560 fold at 10× coverage (**Supplementary Fig. 2**).

#### Error enrichment

The confidence scores of SeqFormer allow most true errors to be isolated in a small candidate subset for downstream correction. Across different platforms and depths, SeqFormer delivers consistently higher F1-scores that take both precision and recall into account than the alignment-based BSAlign (**Fig. 2c, Supplementary Figs. 3**). Using a pragmatic threshold of 0.9 (flagging bases with confidence < 0.9 as candidate erasures), Polus peaks at F1-score with 40.6% under 4× coverage on the Illumina dataset while flagging 0.06% of bases, and 10× coverage with only 0.03% of bases flagged on ONT dataset. Tightening the threshold to 0.8 further improved SeqFormer’s precision without sacrificing recall, yielding F1 up to 42.2% while keeping the flagged set below 0.14% of positions on ONT dataset. These results indicate that SeqFormer provides far stronger error enrichment with orders-of-magnitude smaller flagged bases acrossplatforms and coverages, yielding more informative erasure hints for downstream SDD.

### Encoding module performance

The encoding stage of Polus revealed codec-specific performance trade-offs on sample files **(Fig. 2d)**. For example, the logic density (information bits per nucleotide, i.e., bits/nt) of YYC declines as file size grows because longer indices are needed to address more oligonucleotides, making YYC more efficient for smaller datasets. In contrast, the logic density of Derrick improves with larger files as its fixed matrix padding overhead becomes proportionally smaller, favoring application on larger files. HEDGES and DNA Fountain show nearly size-invariant density. In terms of speed, Derrick is the fastest mainly thanks to the implementation in C language. All codecs produced ∼50% GC-content on average. Fountain, YYC, and HEDGES enforce homopolymer runs≤6 nt, whereas Derrick relies on randomness and occasionally has slightly longer runs. By reporting these metrics, Polus enables informed codec selection based on application-specific priorities such as density, constraint stringency, and speed.

### Multi-stage error simulation

Polus integrates a modular, *in-silico* pipeline that simulates errors arising from synthesis, storage, amplification, and sequencing (**Supplementary Note 2**), enabling real-world DNA storage error landscapes. Unlike prior platforms restricted to Illumina-like substitution errors^24^, Polus extends simulation to both short-read and nanopore long-read profiles by integrating the simulators dt4dds and Badread^30^, respectively. The simulated error statistics match the known platform differences, i.e., under Illumina-like conditions, substitutions dominate (5.2×10^-3^ per base on average) with indels an order of magnitude rarer (mostly single-base events), whereas ONT-like simulations show higher overall error rates (∼5.3%) with indels prevalent (**Fig. 3a, Supplementary Figs. 4-5**).

**Fig. 3.**
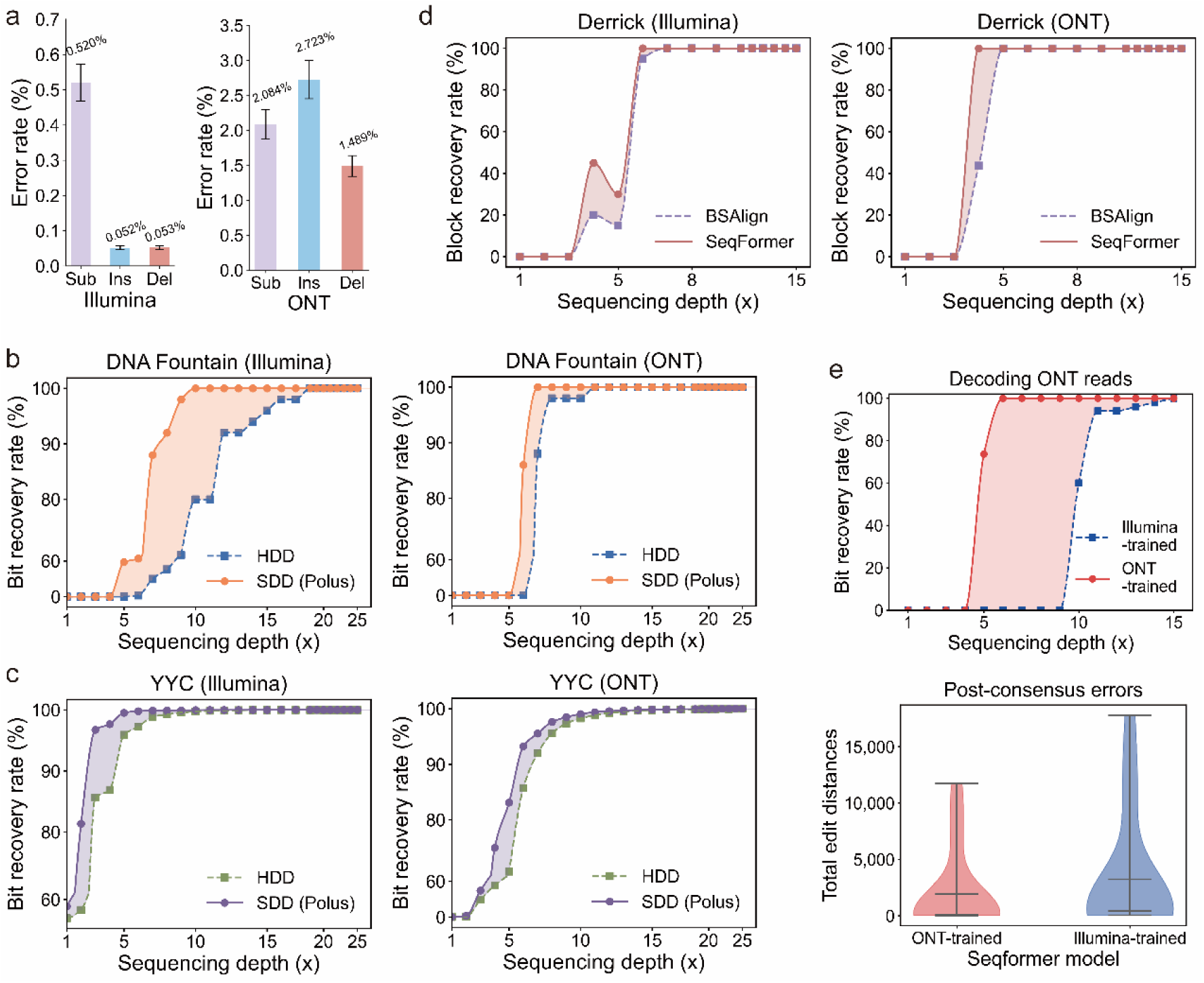
Soft-decision decoding gains under simulated platform-specific error profiles. a, Distribution of error types (substitutions, insertions, and deletions) generated by the Polus simulation module for Illumina and ONT platforms. The simulation reproduces characteristic platform biases, with substitutions dominating Illumina reads and indels prevalent in ONT reads. **b–d,** File recovery rates across sequencing depth for three representative codecs: (b) DNA Fountain, (c) YYC, and (d) Derrick. Dashed lines represent baseline hard-decision decoding (HDD), while solid lines represent Polus-enhanced soft-decision decoding (SDD). Across depths, SDD consistently achieves higher recovery. **(e)** Impact of model specificity on decoding performance. The curve compares the recovery rate of ONT data decoded using a matched ONT-trained SeqFormer model (orange) versus a mismatched Illumina-trained model (grey).

### Decoding module and soft decoding gains

In the Decoding stage, Polus uses the error probability outputs of SeqFormer to implement SDD across different codecs, significantly extending their error**-**correction capability. Rather than treating each base as equally reliable, Polus leverages the per-base posterior probabilities of SeqFormer to mark low-confidence loci as erasures. Since RS codes correct erasures more efficiently than unmarked errors, this approach increases effective ECC headroom without changing the encoding. Consistently, integrating SeqFormer with existing codecs reduced the sequencing depth required for perfect recovery, lowered associated costs, and enhanced the achievable physical storage density in all tested schemes (**Table 1**).

**Table 1.** Overall decoding performance improvement with Polus’ SDD. On *in-silico* Illumina and ONT datasets, soft-decision decoding (SDD; Polus) reduced the quired sequencing depth and increased physical density relative to hard-decision decoders for DNA Fountain, YYC, and BSAlign-guided SDD for Derrick, while eserving perfect file recovery and comparable cost, and runtime. “-” indicates that the hard-decision decoding (HDD) failed to achieve perfect recovery even with ditional sequencing depths.

| Sequencing platform | Codec | Decoder | Required depth (×) | Physical density | File recovery | Cost (\$) | Decoding runtime (s) | Logical density |
| --- | --- | --- | --- | --- | --- | --- | --- | --- |
| Illumina (encoding 108 KB files) | DNA Fountain | SDD (Polus) | 10 | 3.09E+19 | 100% | 97.95 | 0.86 | 1.357 |
|  |  | HDD (baseline) | 18 | 1.72E+19 | 100% | 99.21 | 0.89 |  |
|  | YYC | SDD (Polus) | 16 | 1.94E+19 | 100% | 98.58 | 0.07 | 1.361 |
|  |  | HDD (baseline) | — | — | 99.98% | — | 0.07 |  |
|  | Derrick | SDD (Polus) | 6 | 6.06E+19 | 100% | 1120.68 | 27.96 | 1.593 |
|  |  | SDD (baseline) | 7 | 5.19E+19 | 100% | 1122.5 | 10.4 |  |
| ONT (encoding 108 KB files) | DNA Fountain | SDD (Polus) | 7 | 4.28E+19 | 100% | 517.71 | 1.02 | 1.315 |
|  |  | HDD (baseline) | 11 | 2.73E+19 | 100% | 520.13 | 0.94 |  |
|  | YYC | SDD (Polus) | 24 | 1.35E+19 | 100% | 488.95 | 0.58 | 1.42 |
|  |  | HDD (baseline) | — | — | 99.98% | — | 0.58 |  |
|  | Derrick | SDD (Polus) | 5 | 9.09E+19 | 100% | 1115.01 | 35.85 | 1.593 |
|  |  | SDD (baseline) | 6 | 7.27E+19 | 98.75% | 1116.32 | 17.18 |  |

#### DNA Fountain with SDD

For DNA Fountain codec, SeqFormer-guided SDD markedly lowered the coverage needed for perfect data retrieval under both Illumina and ONT error profiles (**Fig. 3b**). With an inner RS (36, 35) code on Illumina-like data, Polus achieved 100% recovery at 11× coverage, whereas the hard-decision decoder left ∼2% of bits unrecovered at ∼17× coverage and required 20× coverage to fully succeed. Under nanopore-like errors (RS (36, 34)), Polus reached full recovery at 7× coverage versus 11× coverage for hard decoding. At 6× coverage, hard decoding failed entirely while Polus still can reconstructed more than 80% of the files. Thus, context-aware erasures substantially reduced the minimum coverage improved recovery at lower depths.

#### YYC with SDD

For YYC codec, SeqFormer-guided SDD eliminated residual indel errors that the original hard-decoder could not correct **(Fig. 3c)**. Under Illumina-like conditions, two oligonucleotides carrying terminal 2 bp insertions prevented complete recovery even beyond 25× coverage with hard decoding. SeqFormer identified these anomalous insertions with low confidence and marked them as erasures, allowing the outer RS decoder to repair both strands and achieve full retrieval at 20× coverage. Across moderate depths (e.g., 6× coverage), Polus consistently improved recovery rate (to >99% vs ∼97% with hard decoding). Similarly, with ONT-like noise, hard decoding plateaued just below 100% recovery at 25× coverage due to uncorrectable indels, whereas SDD reached 100% recovery at 24× coverage by erasing those errors.

### Derrick with Polus

In the RS (255,223) Derrick scheme, SeqFormer-based decoding reduced the required coverage for error-free recovery **(Fig. 3d)** and improved physical density, while maintaining comparable runtime **(Table 1)**, in contrast to the original BSAlign-guided pipeline. Across all three codecs, these results illustrated that probabilistic soft decoding universally improved reliability, lowered redundancy, and cut down sequencing requirements for error-free recovery.

### Adaptive decoding performance

In practical application, well-calibrated confidence estimates enable an efficient adaptive sequencing strategy to minimize laboratory effort. We simulated this with a DNA fountain dataset (317 oligonucleotides protected by RS (36, 35)). For each starting coverage (5× to 8×), fifty independent resamplings were generated. As a baseline, uniform deepening to 11× coverage was required to achieve 100% recovery with the Polus SDD pipeline. In the adaptive setting (**Supplementary Note 3**), we began at the selected starting coverage, identified decode-failed oligonucleotides, and iteratively requested oligo-specific additional reads. After each increment we recomputed SeqFormer’s estimated consensus accuracy and attempted decoding only when this estimate exceeded 0.998. If decoding fails, another increment was performed. Each oligonucleotide cluster underwent at most five adaptive iterations.

This adaptive strategy consistently achieved 100% file recovery while using orders-of-magnitude fewer additional reads than uniform deepening **(Fig. 4c)**. Across 5× to 8× coverage, the adaptive procedure required 40 to 3,891 additional reads in total, corresponding to 99.92% to 95.91% fewer than the uniform-deepening baseline at the same starting point. Thus, computational guidance via SeqFormer can substitute for brute-force sequencing, preserving data integrity with minimal sequencing overhead.

**Fig. 4.**
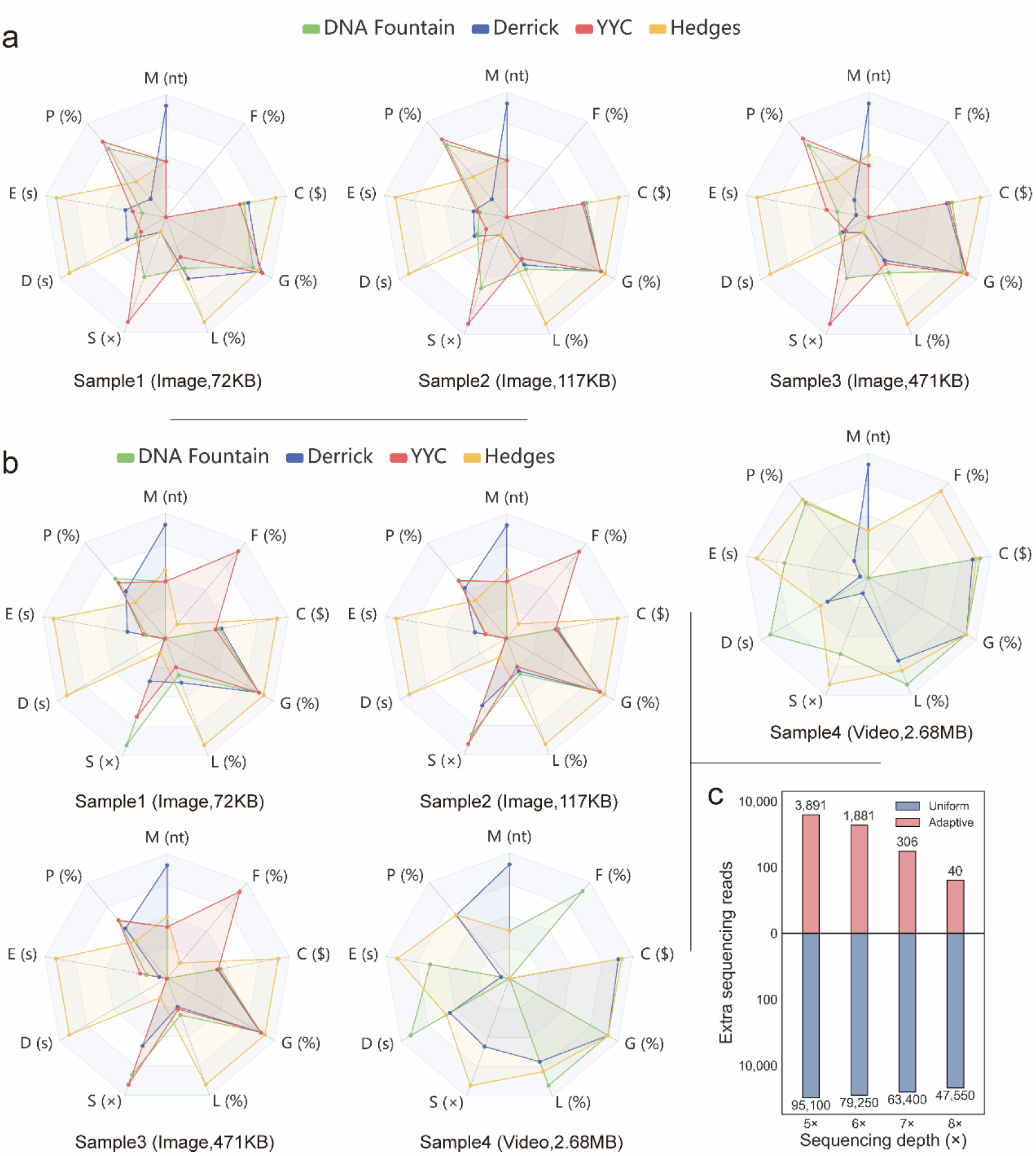
Multi-dimensional performance evaluation and adaptive resequencing efficiency. a-b, Radar charts summarizing the trade-offs of four codecs (DNA Fountain, YYC, HEDGES, Derrick) under (a) Illumina and (b) ONT error profiles. Axes represent nine standardized metrics: File recovery rate (F), Cost (C), Sequencing depth (S), Encoding time (E), Decoding time (D), GC deviation (G), and Max homopolymer length (M), Physical density shortfall (P, defined as the percentage of the theoretical physical storage density (455 EB/g) not achieved by a given scheme, smaller values indicate densities closer to the theoretical limit.), and Logical density shortfall (L, defined as the percentage of the theoretical maximum (2 bits/nt) not achieved.) **c,** Comparison of additional sequencing reads required to achieve full recovery using Polus’s adaptive strategy versus a uniform deepening baseline.

### Platform-specific SeqFormer improves decoding

Tailoring SeqFormer to the sequencing platform further boosted decoding performance **(Fig. 3d)**. We compared two models–one trained on Illumina reads and one on ONT reads–for decoding an ONT test dataset. The ONT-trained model produced ∼40% fewer post-consensus errors on average and achieved full data recovery at 7× coverage, whereas the Illumina-trained model required 15× coverage (a 53.3% higher depth). The results demonstrate that platform-matched training is critical for maintaining calibration and maximizing decoding gains. In practice, we recommend adopting SeqFormer trained on the sequencing technology of interest and fine-tuning on similar error profiles when possible, to fully leverage probabilistic decoding’s advantages.

### Evaluation metrics module

Polus’s evaluation module benchmarked each codec across nine metrics capturing retrieval fidelity, storage density, cost, and biochemical constraints. The metrics are defined detailed in **Methods** section. This multi-dimensional comparison revealed trade-offs that would be obscured by any single-metric assessments. Under Illumina-like conditions, YYC achieved the highest logical density on small files and lowest synthesis cost, but required higher sequencing coverage (lower effective physical density) (**Fig. 4a**). In contrast, Derrick provided better coverage efficiency on small files and dominated large files (≥ 471 KB) with the highest logical density and consequently the greatest physical density. Under ONT-like error conditions, similar trends held. Notably, DNA Fountain failed to recover the largest dataset (∼2.6 MB) due to indel-induced frame shifts that its hard-decision RS decoding could not correct (**Fig. 4b**). HEDGES attained the highest physical density (owing to a minimal 2× coverage requirement) but suffered the lowest logical density and about double the overall cost of other schemes.

Two quantitative trends emerged from cross-platform analysis. First, total costs correlated negatively with logical density, because synthesis dominated expenditure at an approximate 1000:1 cost ratio relative to sequencing^31^. Second, when sequencing depth was incorporated into physical density estimation, the effective physical density was determined primarily by the combined effects of information density and sequencing depth. Because logical density (upper bound ≈2 bits per nt) was less impactful than differences in depth, our results indicated that effective physical density is inversely correlated with sequencing depth. This further implies that, for a given encoding scheme, enhancing the decoder’s error-correction capability not only reduces total cost but also substantially increases physical density.

## Discussion

By learning calibrated per-base error probabilities from multimodal read evidence, Polus converts uncertainty into erasures and feeds RS decoders, consistently reducing the sequencing depth needed for lossless recovery across DNA Fountain, YYC, and Derrick pipelines. To mitigate “false-positive successes” due to short-length RS codes, we further add CRC32 checks in these pipelines which guards against cases where RS decoding passes but the reconstructed content remains incorrect. Benchmark results demonstrate that Polus reduced the coverage required for lossless recovery by 38.9% (DNA Fountain, 18×→11×) and eliminated systematic indel errors in YYC schemes (BER 0.03%→0).

The decoding gains are primarily driven by SeqFormer, which integrates multi-modal channel signals within a context-aware Transformer to produce calibrated base-wise error probabilities. Ambiguous loci are then erased for RS decoding, effectively expanding the correctable set without increasing parity. Importantly, certain errors—especially indels in DNA Fountain workflows—do not vanish with depth alone; soft-decision inputs, however, enable the outer code to reconcile these events without extra redundancy, which explains the observed reduction in coverage thresholds for zero-loss recovery. SeqFormer can also generate high-confidence consensus sequences from multiple reads of the same locus, suggesting its potential utility as a tool for multiple sequence alignment and related bioinformatics applications.

Beyond decoding, Polus provides a standardized evaluation suite and a multi-stage channel simulator, thereby complementing earlier frameworks that primarily emphasized codec aggregation. Our cross-codec benchmarks provide practical guidance: for applications prioritizing cost and effective physical density. Derrick offers the most balanced performance across platforms and scales better as file size grows. YYC delivers higher logical density for small files but, lacking explicit indel handling, benefits from SeqFormer-guided soft-decision decoding to achieve stable recovery. DNA Fountain, Derrick, and HEDGES show robust recovery under moderate error conditions. Taken together, these observations argue for probabilistic, confidence-guided decoding as a default design choice, with the codec selected according to file size and deployment constraints. We recommend adoption of this evaluation suite as a benchmark so that future DNA storage studies are comparable and reproducible across codecs, chemistries, and datasets. The platform’s modular design further enables seamless integration of both established and newly developed codecs; for instance, future codecs can be coupled with the Polus error predictor and a soft-input decoder to form a context-aware “soft” decoding pipeline.

Looking ahead, an important direction is to extend SeqFormer to expanded molecular alphabets, particularly composite letters, which suffer from elevated error rates and exhibit strong letter-specific transition biases. By exploiting cross-read contextual correlations, SeqFormer could more accurately calibrate transition tendencies among easily confusable letters, thereby improving inference fidelity. In parallel, integrating SeqFormer’s context-aware posterior probabilities with composite-letter priors (transition libraries) would enable prior-guided re-ranking of candidate letters, effectively reconciling probabilistic predictions with the molecular error profiles arising from synthesis and sequencing. Taken together, these strategies outline a principled route to reduce sequencing depth requirements and enhance robustness in composite letter-based DNA storage systems. Beyond algorithmic advances, integrating Polus with automation-ready laboratory workflows would further allow coding decisions (e.g., re-write versus re-read) to be guided by calibrated uncertainty in real time. Collectively, these steps can move DNA storage toward lower cost and reliability-aware operation, while preserving codec diversity.

## Methods

### SeqFormer model

#### Model architecture

SeqFormer is a Transformer-based model designed to infer a per-base consensus sequence and a calibrated error probability for each position from a cluster of sequencing reads corresponding to one DNA oligonucleotide. For each oligo’s read cluster (with 𝑁 reads and aligned length 𝐿), we construct two input tensors: (1) an aligned nucleotide matrix 𝑋_𝑠𝑒𝑞_ ∈ 𝛴^𝑁×𝐿^ (Σ={A, C, G, T, -}, where “-” denotes a gap in the multiple sequence alignment); (2) an aligned Phred-quality matrix 𝑋_𝑞𝑢𝑎𝑙_ ∈ 𝑄^𝑁×𝐿^ (per-base Phred quality scores from FASTQ files, 𝑄 represents the set of ASCII characters used to encode quality values). Each branch is passed through a learned embedding layer that projects the inputs to a fixed-dimensional vector (we use an embedding dimension 𝑑 = 512), followed by a root-mean-square layer normalization (RMSNorm) to stabilize training across varying coverages. The per-read features from the nucleotide branch and quality branch are concatenated along the feature dimension and fused using a 1D pointwise convolution layer (kernel size = 1), yielding per-read representations 𝐻 ∈ 𝑅^𝑁×𝐿×𝑑^. We then aggregate information by sum-pooling over reads to obtain a single fused representation 𝑍 ∈ 𝑅^𝐿×𝑑^. This aggregated sequence 𝑍 is fed into a Transformer encoder for contextual modeling. The Transformer encoder comprises 8 encoder layers, each integrating the multi-head self-attention module (8 heads) and the feed-forward module. Both submodules employ RMSNorm-based pre-normalization and residual connections. The encoder outputs a contextual embedding 𝑍~ ∈ 𝑅^𝐿×𝑑^ that integrates long-range and cross-context dependencies that simple majority voting may miss. Finally, a linear projection followed by a softmax activation yields a probability distribution over {A, C, G, T, -} at each position. The base with the highest probability is taken as the consensus call for that position, and the softmax probability of that consensus base serves as the model’s confidence score (i.e., the estimated probability that the call is correct).

#### Model training

We trained SeqFormer by minimizing the per-position cross-entropy between its predicted categorical distribution over {A, C, G, T, ‘–’} and the known designed oligo reference aligned to each read cluster. The model was implemented in PyTorch and optimized using the Adam optimizer (initial learning rate 0.0002, *β*₁=0.9, *β*₂=0.999). We applied a weight decay of 0.01 and dropout of 0.1 in the Transformer layers to regularize the model. Training was run for up to 5 epochs with early stopping based on validation loss to prevent overfitting. We selected the model checkpoint with the highest validation accuracy (consensus bases matching the ground truth) and best calibration (lowest deviance between predicted confidence and actual error rate). All training and inference were accelerated using an NVIDIA RTX A6000 GPU.

#### Datasets and preprocessing

Because DNA sequencing error profiles differ by platform, we trained platform-specific instances of SeqFormer with identical architecture but separate weights for Illumina short reads and ONT long reads. Training used previously published DNA-storage datasets: Illumina dataset from Organick *et al*.^17^ encoding ∼200 MB and ONT dataset from Ding *et al.*^13^ encoding ∼5 MB. Each dataset was split 3:1 into training and test sets. Accession codes and download links are provided in Supplementary Table S1. For each dataset, raw FASTQ files were demultiplexed and adapter-trimmed, then clustered by oligonucleotide address to form per-oligo read piles. Each pile was aligned using BSAlign to generate a multiple sequence alignment, with gaps represented as “-”; the corresponding per-base Phred quality scores were retained and mapped to the alignment coordinates.

#### Model selection

SeqFormer is distributed as a single executable with a command-line option to select the pretrained model: ONT-trained is invoked with “SeqFormer --tgs”, and Illumina-trained with “SeqFormer --ngs”. Unless specified otherwise, we used frozen weights without additional fine-tuning.

### SDD implementation in Polus

After SeqFormer produces a consensus sequence and an error probability for each base of an oligo, we then feed this information into the decoding algorithms of Fountain, YYC, or Derrick as appropriate, modifying each decoder to accept “soft” inputs. Across all three pipelines, soft information is ultimately consumed at an RS layer.

#### Principle

An RS code over 𝑛 symbols (with 𝑘 data symbols) can correct up to 𝑡 = [(𝑛 − 𝑘)/2] errors (or 2t erasures), and more generally can successfully decode if the condition 2𝑒 + 𝑠 ≤ (𝑛 − 𝑘) is met (where 𝑠 is the number of erasures supplied to the decoder and 𝑒 is the number of residual errors). We leveraged SeqFormer’s basewise confidence to identify likely error positions in the consensus. Specifically, we define a confidence threshold 𝜃 (by default 𝜃 = 0.1) on SeqFormer’s predicted error probabilities. Any base with 𝑃_𝑒𝑟𝑟𝑜𝑟_ > 𝜃 is marked as a candidate erasure. These candidates are ranked by their error probability to form an erasure set 𝐸. We then iteratively select subsets of 𝐸 for RS decoding, gradually pruning the least likely erasures until a valid codeword is obtained. Successful decodings are verified against a CRC32 checksum, which ensures that undetected RS collisions cannot propagate as false positives.

#### Threshold and calibration

SeqFormer’s confidences are strongly polarized in practice (correct bases typically > 0.9). A sweep of 𝜃 ∈ [0, 0.2] shows a broad optimum around 𝜃 = 0.1, which maximizes full-file recovery by balancing false erasures against missed errors. We set 𝜃 = 0.1 as default.

#### Codec-specific mapping

In DNA Fountain, base-level confidences within each droplet are aggregated to byte-level bytes exceeding 𝜃 are erased before inner RS decoding, and only RS-passed droplets are forwarded to LT solving. In YYC, the outer RS code spans strands; per-strand symbols inherit from their protected segments, and soft erasures allow the parity code to focus capacity on genuinely ambiguous symbols. In Derrick, fixed-length payload sequences form a matrix; RS is applied across sequences (columns). For each column, symbol probability comes from the aligned base window of that column; erasures are flagged column-wise and decoded vertically.

### Evaluation metrics

To compare performance across schemes comprehensively, we defined nine key metrics in Polus as follows:

#### File recovery rate

This category includes: Bit recovery rate (BRR)–the fraction of bits that are correct in the final reconstructed file (after decoding) relative to the original input–and Block recovery rate (LRR)–the percentage of codeword blocks that are decoded without any errors. In our context, a 100% BRR or LRR means the file is perfectly decoded.

#### Logical density

The number of information bits stored per nucleotide synthesized. Let 𝐵_𝑝𝑎𝑦𝑙𝑜𝑎𝑑_ be the number of information bits in the input file(s), and let 𝑁_𝑛𝑡_ be the total number of synthesized nucleotides (sum over all oligos, including addressing and ECC). Logical density is

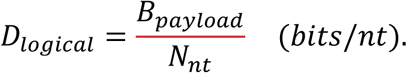

#### Physical density

The amount of data stored per unit mass of DNA, typically expressed in bytes per gram. Firstly, under an ideal dry-DNA assumption, the number of nucleotides per gram is 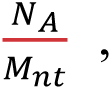 where 𝑁_𝐴_ = 6.02 × 10^23^ 𝑚𝑜𝑙^−1^ is the SI-defining Avogadro constant and 𝑀_𝑛𝑡_ is the average molar mass per nucleotide of ssDNA (we use the standard approximation 𝑀_𝑛𝑡_ = 330 𝑔 · 𝑚𝑜𝑙^−1^ · 𝑛𝑡^−1^). Thus, the ideal physical density is computed by

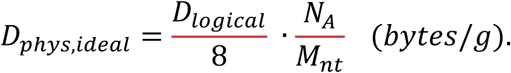

Then actual physical density for ssDNA. In practice, reliable readout requires multiple copy numbers per oligo, 𝐶, which we take as the minimal average coverage that yielded error-free file recovery in our pipeline (from our *in-silico* experiments). Under the assumptions the storage of 100 molecules per oligo sequence^14^, no synthesis/long-term loss (effective yield 𝑓_𝑦𝑖𝑒𝑙𝑑_ = 1), the actual density reduces to 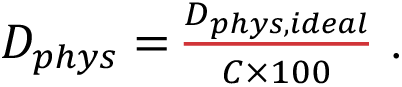 More details about the computation were added in the Supplementary note 4.

#### Sequencing depth requirement

The minimum average sequencing coverage per oligo required to consistently recover the data without errors. We determine this by running simulations with decreasing read depths until decoding fails.

#### Encoding runtime

We measured wall-clock times of the encoder implementations on an Intel Xeon Platinum 8352V processor to compare their efficiency.

#### Decoding runtime

We measured wall-clock times of the decoder implementations on an Intel Xeon Platinum 8352V processor to compare their efficiency. The clustering, SeqFormer inference (for Polus), and error correction decoding time were separately recorded.

#### Costs

An estimate of the monetary cost to store and retrieve the data using each scheme, factoring in DNA synthesis cost (per base, given the logical redundancy) and sequencing cost (per read, given the depth needed). We used current synthesis prices (∼$0.10 per base) and sequencing prices ($1000 per ∼1 billion reads for Illumina, ∼$30 per GB for Nanopore MinION) as a basis^32^.

#### GC contents

The average (and distribution of) GC fraction in the encoded oligos. All our schemes enforced roughly 45–55% GC content, but we included this to ensure none produce extreme GC outliers that could impede PCR or sequencing.

#### Maximum homopolymers length

The longest run of identical nucleotides in any encoded oligo. This is a measure of biochemical constraint adherence, as long homopolymers (e.g., >5 bases) can cause higher sequencing errors.

## Data availability

All the datasets used for training and evaluation were obtained from public databases. The Illumina sequencing datasets from Organick *et al.*^17^ and the Nanopore sequencing datasets from Ding *et al.*^13^ used to train SeqFormer can be obtained via their supplementary materials; we provide processed versions (read alignments and per-base data) in our repository to facilitate model training. The accession numbers and data links are listed in Supplementary Table 1. All simulated data generated during this study (including encoded oligo libraries, simulated FASTQ reads, and decoder outputs) can be regenerated using the code and parameter settings we have released. The sample image and video Files for Polus pipeline tests, decoding performance and evaluation performance are provided at https://github.com/dinglulu/Polus/tree/main/test_data.

## Code availability

All source code of the Polus platform, including the SeqFormer implementation, codec algorithms, the simulation pipeline, and the soft-decision decoding and evaluation module, is available in an open-source repository at https://github.com/dinglulu/Polus. The repository provides documentation and scripts to reproduce all experiments reported in this work. A web-hosted instance of Polus is available at https://polus.bioailab.net/polls/home. The SeqFormer model is also released as a standalone repository at https://github.com/dinglulu/SeqFormer.

## Author Contributions

L.D. and Z.Z. conceived the idea of Polus. Z.Z., and L.L. coordinated and supervised the project. L.D. and S.X. designed the SeqFormer algorithm, S.X., H.Z. and J.W. developed and implemented the algorithm. L.D. Z.Z. and L.L. designed the Polus platform, K.W. and H.Z. developed the platform, performed experiments, and analyzed data. L.D., Z.Z. and L.L. drafted the manuscript. W.Z., B.L., and G.W. provided critical suggestions on algorithm evaluations and improved the manuscript. All authors discussed the results and approved the final manuscript.

## Declaration of Interests

The authors declare no competing interests.

## Acknowledgements

This study was supported by the National Key Research and Development Program of China (2022YFF1202104) and National Natural Science Foundation of China (32401256).

## Notes

### Competing Interest Statement

The authors have declared no competing interest.

