## Supplementary Material for "Polus: a Transformer-based Soft-decision Codec Enhancement Platform for DNA Storage"

**This PDF file includes:**

Supplementary Notes 1– 4
Supplementary Figures 1– 5
Supplementary Table1

### Supplementary Notes

#### Supplementary Note 1: Implementation details of Polus encoding schemes and decoding experiments

##### Polus encoding schemes.

**Integration of multiple DNA codecs:** The Polus platform employs a modular plugin interface to support diverse DNA encoding/decoding architectures. In this study, we integrated four representative codecs—DNA Fountain, Yin–Yang Codec (YYC), HEDGES, and Derrick—by reimplementing them according to their original specifications and encapsulating them within the Polus API for seamless interoperability. All schemes convert binary input files into a library of DNA oligonucleotide sequences containing necessary addressing indices and error-correction redundancy.

**DNA Fountain**: We implemented DNA Fountain following Erlich *et al.*[1]. In our configuration, input data was chopped into fragments which were encoded into "droplets" of a predetermined payload size; an RS error-correcting code was applied as an inner code to each droplet (adding parity nucleotides within the droplet), and a Luby Transform (LT) erasure code was used as an outer code across droplets. The logical redundancy was set to approximately 12% (determined by the ratio of generated droplets to the source data length).

**YYC**: We implemented the Yin–Yang encoding based on Ping *et al.*[2]. YYC maps pairs of binary bits to single nucleotides using context-dependent "Yin" and "Yang" rules to satisfy biochemical constraints. This yields a theoretical coding rate of 2 bits per nucleotide (nt). To ensure robustness, we appended an outer RS (36, 34) code, introducing 2 parity symbols for every 34 data symbols.

**HEDGES:** We integrated HEDGES following the procedure outlined by Press *et al.*[3]. Data bits are encoded into DNA sequences utilizing synchronization markers (hash anchors) and convolutional parity segments to correct insertions, deletions (indels), and substitutions. We configured HEDGES to maintain a total redundancy of 10–15%.

Output sequences were truncated or padded to a standard length of 150 nt and verified against GC content and homopolymer constraints.

**Derrick**: Implemented the matrix-coded architecture described by Ding *et al.*[4], featuring a vertical RS outer code optimized for soft-decision decoding. The bitstream is segmented into fixed-length payload sequences, upon which an RS code is applied column-wise. We utilized an RS (255, 223) code over the Galois Field $GF(2^8)$, generating 32 parity bytes per column.

Each encoder output a FASTA library of oligonucleotides. Polus automatically compiles an encoding report quantifying the library's characteristics, including logical density, encoding runtime, and biochemical feature distributions (GC content and homopolymer run lengths).

**Decoding experiments implements.**

We evaluated Polus's soft-decision decoding capabilities across three DNA storage codecs under *in-silico* experimental conditions. For DNA Fountain and YYC, we benchmarked Polus (SeqFormer-guided SDD) against baseline hard-decision decoders (HDD). For Derrick, we compared the Polus pipeline (SeqFormer-driven) against the baseline BSAlign-based soft decoder.

**Code Parameters:** DNA Fountain and YYC employed short RS codes over $GF(2^8)$ with small parity budgets ($n - k = 1\ or\ 2$; inner for Fountain, outer for YYC), while Derrick used a longer RS (255, 223).

**Error Profiles:** All codecs were evaluated under two distinct error conditions simulating Illumina (short-read) and ONT (long-read) sequencing platforms, generated via standard simulation pipelines.

#### Supplementary Note 2: *In silico* DNA channel simulation

To rigorously benchmark decoding performance without the prohibitively high cost of wet-lab experiments, we implemented a comprehensive in silico simulation pipeline. This modular simulator models the stochastic errors accumulated across the entire DNA data storage lifecycle: synthesis, storage (aging), PCR amplification, and sequencing.

**Synthesis errors:** Oligonucleotide synthesis was modeled using *dt4dds*[5], a stochastic simulator that introduces context-dependent substitutions, insertions, and deletions characteristic of phosphoramidite chemistry.

**Storage and decay:** We simulated DNA degradation to model varying storage durations. The simulator accounts for random strand breakage and base damage events (e.g., Cytosine deamination to Uracil/Thymine). For accelerated aging benchmarks, damage rates were elevated to emulate multi-year storage.

**PCR amplification:** To mimic library preparation, we implemented a simplified PCR model accounting for bias and polymerase errors. Each oligonucleotide was assigned a coverage multiplier drawn from a distribution with a dispersion factor reflecting empirical PCR bias, simulating the over- or under-representation of specific strands. Polymerase-induced errors (substitutions and indels) were also superimposed.

**Sequencing errors:** Polus integrates platform-specific sequencing simulators. For Illumina sequencing, we utilized *dt4dds* in sequencing mode to generate reads with predominantly substitution errors (e.g., 0.1% per base) and minor indel rates. For ONT (long-read) sequencing, we leveraged the read simulator *Badread*[6] to generate long reads exhibiting typical nanopore error profiles, characterized by higher total error rates (~5%) and a prevalence of indels.

The final output is a set of FASTQ files mimicking raw experimental data.

**Supplementary Note 3: Adaptive decoding for *in silico* experiments.**

We assessed the efficiency of Polus's adaptive decoding using a DNA Fountain dataset comprising 317 oligonucleotides protected by an RS (36, 35) code. For each initial coverage level $\alpha \in (5\times,\ 6\times,\ 7\times,\ 8\times)$, we performed fifty independent resampling using the same upstream processing and decoder configuration as in the main pipeline.

**Uniform Deepening Baseline:** We first established the minimum coverage required for complete recovery using a uniform deepening approach. Starting from 5×, coverage was increased stepwise until the file was 100% recovered at 11×. The total additional read count for the uniform baseline from a starting coverage $\alpha$ is calculated as:

$$\text{Uniform additional reads } = (11 - \alpha) \times 317 \times 50.$$

**Adaptive Resequencing Strategy:** Instead of uniformly sequencing all strands, the adaptive strategy identifies specific oligonucleotides that failed decoding at the initial coverage. Additional reads were requested for these targets in small, fixed batches. After each batch, we recomputed the consensus accuracy estimated by SeqFormer.

**Termination Criteria:** Decoding is attempted only when SeqFormer's estimated confidence for a target oligonucleotide exceeds a threshold of 0.998.

**Loop:** Successful decoding terminates the process for that oligonucleotide. If decoding fails or confidence remains low, another batch of reads is requested, up to a maximum of five adaptive iterations.

The proportion of reads saved relative to uniform deepening at the same starting point, computed as

$$Reads\ saved\ = 1 - \frac{\text{Adaptive additional reads}}{\text{Uniform additional reads}}.$$

**Supplementary Note 4: Definition of the actual physical density.**

**Actual physical density** ($D_{phys}$) represents the information storage capacity per unit mass of DNA, is computed in three steps:

(1) Compute $D_{logical}$ from the encoded FASTA by counting $N_{nt}$ and dividing the known payload bits $B_{payload}$ .

(2) Covert $D_{logical}$ to $D_{phys,ideal}$ using $N_A$ and $M_{nt}$.

(3) Determine $C$ from decoding experiments.

Notes: If future experiments include duplex storage or explicit yield losses, the general form $D_{phys} = \frac{D_{phys,ideal}}{C} \times f_{yield} \times \frac{1}{f_{ds}}$

Should be used, where $f_{ds} = 2$ for dsDNA. For intuition, the classical theoretical upper bound[7] of $455\ EB/g$ is recovered by setting $D_{logical} = 2, C = 1, and\ M_{nt} = 330\ g/mol$.

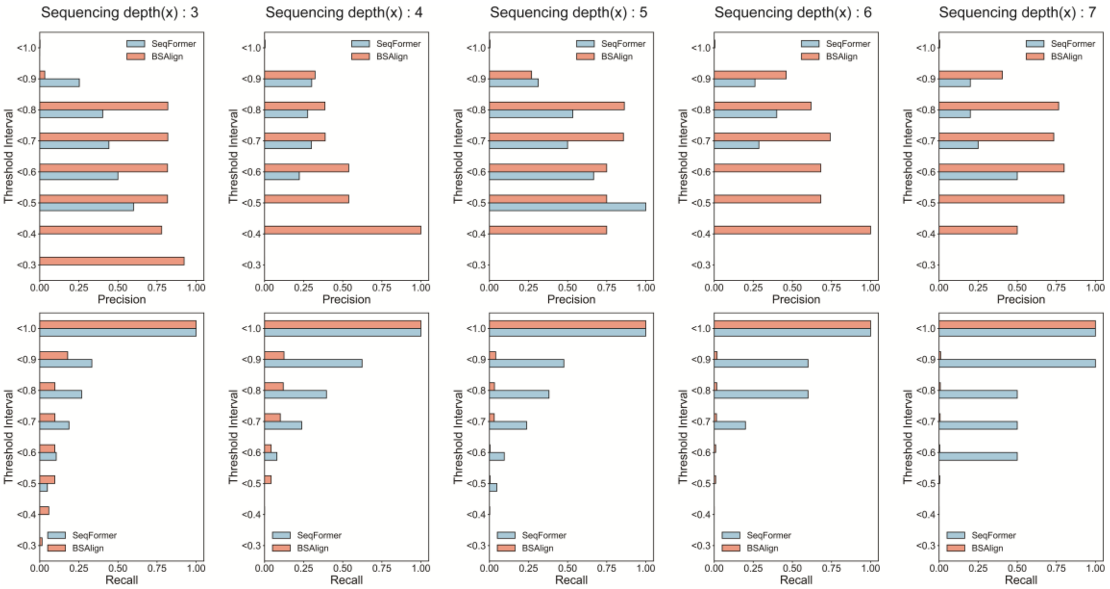


**Supplementary Figure 1. SeqFormer's confidence calibration on the Illumina dataset.** We stratified nucleotide's positions into confidence bins (y-axis) based on the model's output probabilities. For each bin, we report Precision (fraction of flagged positions that are true errors) and Recall (fraction of all true errors captured within the flagged set) across varying sequencing depths (3× to 7×).

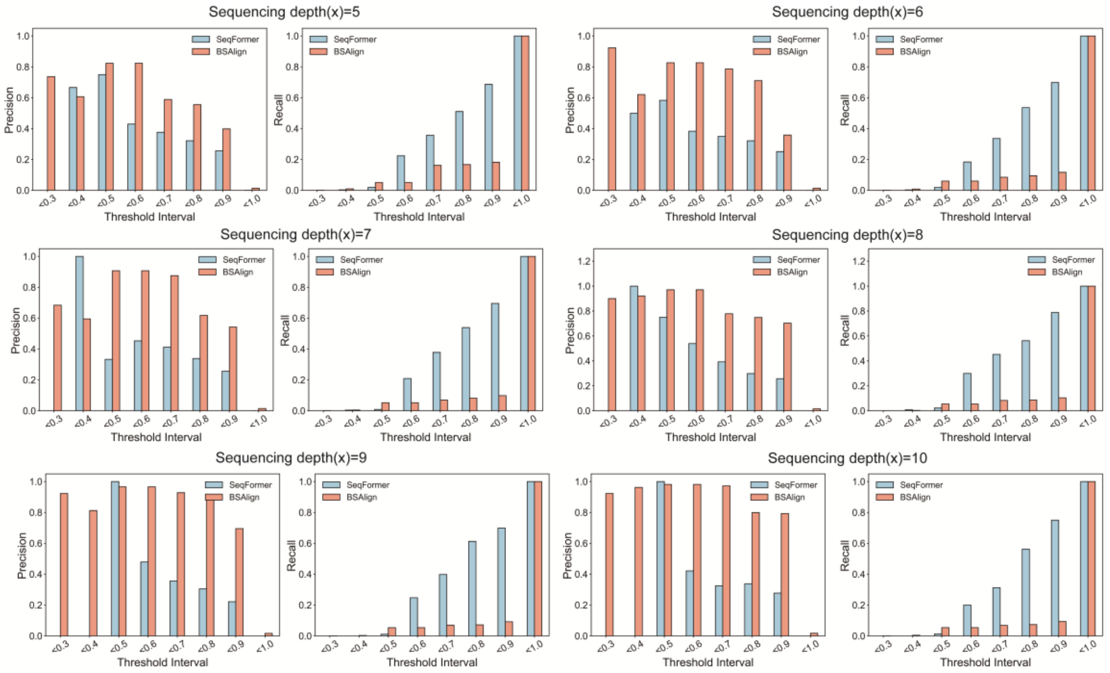


**Supplementary Figure 2. SeqFormer's confidence calibration on ONT dataset.** Similar to Supplementary Fig. 1, this figure illustrates the model's calibration performance across sequencing depths from 5× to 10×.

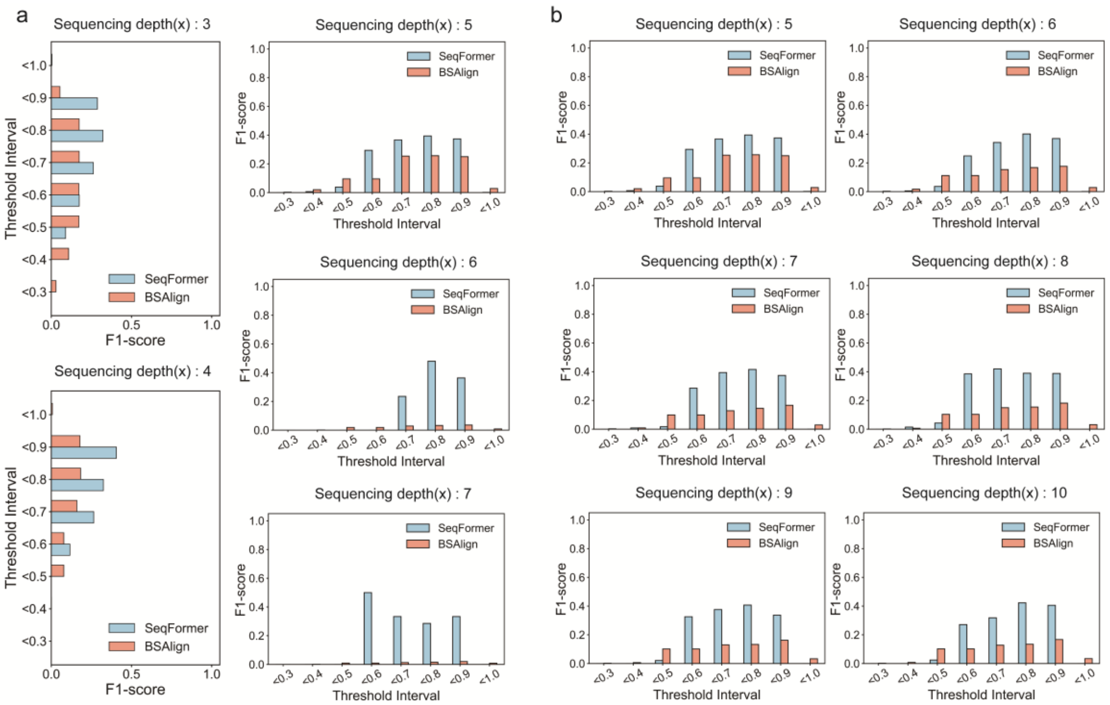


**Supplementary Figure 3. Comparison of SeqFormer and BSAlign in calibrated error enrichment.** Evaluated on the identical Illumina (a) and ONT (b) datasets characterized in Supplementary Figs. 1 and 2, this figure extends the analysis to quantify the overall efficiency of error identification, utilizes the F1-score (the harmonic mean of precision and recall) as a comprehensive statistic to benchmark detection performance.

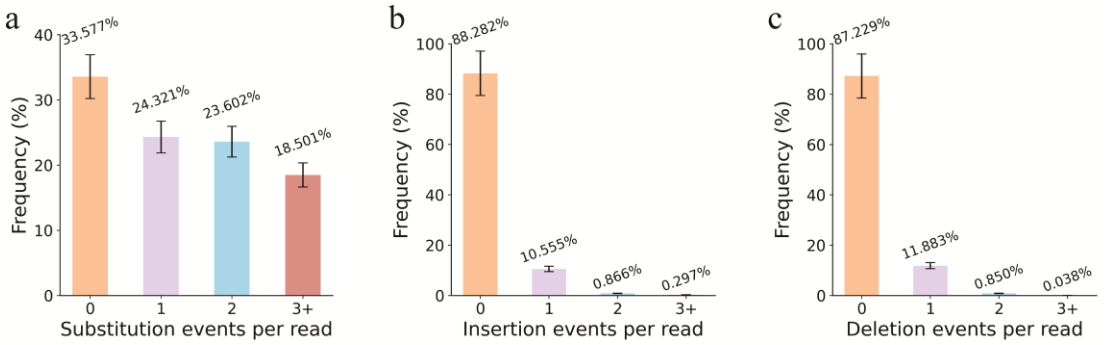


**Supplementary Figure 4. Error type distribution under Illumina-like simulation. a-c,** Frequency distribution of error events per read for substitution (a), insertions (b), and deletions (c). The simulation reflects the characteristic error profile of Illumina platforms, where substitution errors are predominant (occurring frequently per read), while insertions and deletions are rare, typically appearing as single-nucleotide events.

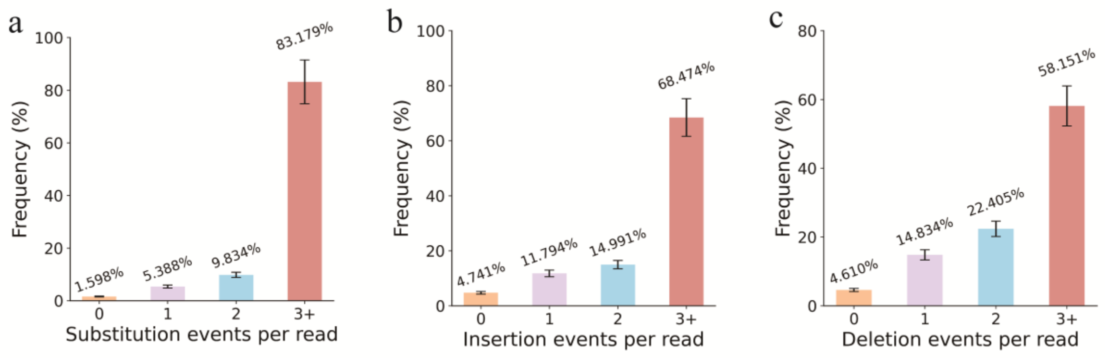


**Supplementary Figure 5. Error type distribution under ONT-like simulation.** a-c, Frequency distribution of error events per read for substitution (a), insertions (b), and deletions (c) under Nanopore conditions. In contrast to Illumina, the ONT profile is characterized by a higher overall error rate and a significant prevalence of multi-nucleotide insertion and deletion events (indels).

**Supplementary Table 1. Datasets used for SeqFormer training and evaluation.** Sources and accession codes for the public datasets used to train the platform-specific SeqFormer models.

| Sources | data links or accession number |
|---|---|
| Organick *et al.*[1] | https://github.com/uwmisl/data-nbt17/tree/master |
| Ding *et al.*[4] | accession number CRA008036 under CNCB Genome Sequence Archive (GSA) |
